# Insights into the genomic features, lifestyle and therapeutic potential of B1 sub-cluster mycobacteriophages

**DOI:** 10.1101/2023.05.30.542743

**Authors:** Ritam Das, Ritu Arora, Kanika Nadar, Saroj Saroj, Amit Kumar Singh, Shripad A Patil, Sunil Kumar Raman, Amit Misra, Urmi Bajpai

## Abstract

**Background:** A large number (about 1200) of mycobacteriophages (phages) have been isolated on *Mycobacterium smegmatis* mc^2^155. Their genome analysis shows high sequence diversity; therefore, based on nucleotide sequence similarity and genomic architecture, the related phages have been grouped in clusters and sub-clusters. However, a deeper study of mycobacteriophages has been conducted only for a few clusters. This study explores the traits of phages belonging to the B1 sub-cluster. We have attempted to functionally annotate and experimentally characterize B1 phages to get an insight into their biology and explore their therapeutic potential.

**Methods:** Analysis of B1 sub-cluster phage genomes to understand their key characteristics & lifestyle and to determine the putative function of hypothetical proteins (HPs), we developed a framework with a specific set of computational tools available online. For the experimental characterization, mycobacteriophages were isolated from environmental samples and were examined for their morphology, lysogeny status, effect on biofilm and activity against drug-resistant *M. smegmatis*. The B1 sub-cluster phages were identified by PCR using the specific primers.

**Results:** We have predicted the function of about 55% of the 77 representative proteins in B1 phages, which were previously deemed hypothetical. We studied ten B1 phages (*Phages 1-10)* which included their morphological characteristics, lysogeny status and antibiofilm activity. TEM analysis, showing an average head & tail size of 65 nm and 202.12 nm, respectively. The turbid morphology of several plaques suggested these phages to be temperate. To verify, we tested their potential to lysogenize *M. smegmatis* and later found the spontaneous release from the putative lysogens. Interestingly, a putative RepA-like protein was identified in B1 phage genomes, indicating a possibility of extrachromosomal replication of prophages. Further, the impact of *Phages 1-10* on *M. smegmatis* biofilm was found to be potent; the highest inhibitory and disruptive effect of phages (at a fixed titre of 10^8^ pfu/ml) was 64% and 46%, respectively. Also, all ten phages could kill 4XR1 (the isoniazid-resistant *M. smegmatis* strain).

**Conclusion:** We believe this combination of experimental analysis and exploration of genomic features of mycobacteriophages belonging to a sub-cluster can provide deeper insights into mycobacteriophage biology and also help in understanding their therapeutic potential.

## 1. Introduction

Mycobacteriophages are phages that infect *Mycobacterium spp.* As per the Actinobacteriophage database, there are over 12,580 mycobacteriophages discovered from the environment, of which the genome of 2257 mycobacteriophages has been completely sequenced (May 2023) (https://phagesdb.org/). Mycobacteriophages show a high degree of genome sequence diversity and mosaicism and possess many uncharacterized genes. The classification of mycobacteriophages into clusters and sub-clusters is based on their overall nucleotide similarities and differences. Phage genome analysis sheds light on viral diversity and evolution and provides genetic information to better understand the mycobacterial host (1). A steady rise in reported cases of antibiotic resistance has prompted a renewed interest in phage therapy (2, 3). Using phages in therapy primarily focuses on their isolation and characterization followed by *in vitro* and *in vivo* screening against drug-resistant bacterial strains. Phage genes encoding undesirable proteins, such as those coding for antibiotics resistance, lysogeny or virulence factors, should be edited before they are considered for therapy. Therefore, genomic, and experimental characterization of phages is essential for their safe and effective use as therapeutics.

The Mycobacteria spp are wearisome to be treated efficiently with antibiotics alone (4); their biofilm-forming ability further adds to the tolerance against antibiotics and other possible treatments (5). Biofilms pose a considerable challenge in treating infectious diseases, as they act as a cell reservoir at the site of infection to repopulate it as soon as the treatment is withdrawn, causing the recurrence of disease (6). Bacterial biofilms also protect the pathogen from the host immune system, resulting in treatment failures and thus causing severe chronic infections (7). The abundant presence of phages in nature and their antibiofilm activity make them valuable candidates as antibacterial agents (8). However, therapeutic consideration cannot be approached without gaining insights into the genomic characteristics of bacteriophages (9).

Currently, there are 32 established mycobacteriophage clusters (A to Z, AA to AE), with the cluster A being the largest, followed by B; and 7 are singletons genetically unrelated to the other clusters. Of the 32 clusters, 12 clusters are further classified as sub-cluster. Cluster B comprises 388 mycobacteriophages (as of May 2023) and is divided into 13 sub-clusters (B1-B13) (https://phagesdb.org/), B1 being the highest populated sub-cluster with 254 phages. Using a combination of *in silico* tools available for genome annotation, we have tried to assign putative functions to several of the hitherto hypothetical genes in B1 phage genomes.

While isolating mycobacteriophages from the soil samples collected from different regions in and around Delhi, India, using *M. smegmatis* (mc^2^155) as the bacterial host, we found B1 phages to be more in number (a frequency of 1 in 3). Information on B1 phages is available in a few earlier reports (10–13), however a detailed insight into their genome and experimental data is limited. In this study, 30 new mycobacteriophages were isolated from 160 soil samples and were screened by a PCR-based method (14) for cluster classification, where primers specific to a conserved region in the tape measure protein of B1 sub-cluster phages were used. Out of thirty, ten mycobacteriophages were found to belong to B1 sub-cluster, which were further analysed experimentally.

## 2. Methods and Materials

### 2.1 In silico analysis of phage genomes

Genome annotation of representative B1 sub-cluster phages (Aelin, GeneCoco, Ashraf, and Maskar) and their comparison were made using DNA Master (http://cobamide2.bio.pitt.edu/) and Phamerator (15), respectively. ARAGORN (16) was used for detecting tRNA and SigA-like promoter regions in the phage genomes and was identified using the built-in feature in DNA master (http://cobamide2.bio.pitt.edu/). Homology analysis was done using BLASTp (17) and HMMER (18). Structural and membrane proteins were identified through PhANNs (19) and TMHMM (20), respectively. The tool PFP-FunDSeqE (21) was used to detect fold patterns. ToxDL (22) and DP-Bind (23) were used to detect the toxicity and DNA binding proteins, respectively. To predict genes encoding depolymerase enzyme in the minor tail protein of B1 sub-cluster mycobacteriophages, sequences of structural proteins were downloaded from UniProt (https://www.uniprot.org/) and analyzed using the PHYPred tool (24). The structural proteins predicted as ‘enzymes’ were then analyzed using I-TASSER (25).

### 2.2 Bacterial strains and growth media

The non-pathogenic and fast-growing *Mycobacterium smegmatis* strain mc^2^155 was used as a host for phage isolation. This strain was grown for 48 hours at 37°C using Middlebrook 7H9 broth containing Carbenicillin (50 μg/ml) and Cycloheximide (10 μg/ml) (HiMedia, India), supplemented with 0.4% glycerol. To check the susceptibility of phages to a drug-resistant strain of *Mycobacterium smegmatis* [Isoniazid (INZ)-resistant strain (4XR1)] and to a NTM [*Mycobacterium fortuitum* (NIHJ1615)], Middlebrook 7H9 broth was used without the Carbenicillin*. M. smegmatis* 4XR1 strain was cultured with *INZ at a* concentration of 40 μg/mL. Susceptibility to *Mycobacterium tuberculosis* (H37Rv) was carried out at the National JALMA Institute for Leprosy & Other Mycobacterial Diseases, India according to a previously published protocol (26).

### 2.3 Isolation, purification, and amplification of mycobacteriophages

Briefly, soil samples were collected from 158 different regions in Delhi-NCR, India. To 25 grams of each sample, an equal volume of phage buffer (100 mM Tris pH 7.5, 100 mM MgSO_4_, 1 mM CaCl_2_, 68 mM NaCl) was mixed and kept undisturbed. The buffer in the top layer was collected, centrifuged, and the supernatant was sterilized using 0.45 μm syringe filters. For enrichment, 2% LB media (Himedia, India) and *M. smegmatis* mc^2^155 culture was added to the filtrate and incubated overnight at 37 °C at 100 rpm, followed by centrifugation at 8,000 rpm. The supernatant obtained was sterilized using a 0.2 μm syringe filter and tested for the presence of phages by the double agar overlay method (12). Phage plaques observed after 48 hours of incubation at 37 °C were purified (with three rounds of purification) and amplified to a phage titre of 10^10^-10^12^ pfu/ml and stored.

### 2.4 Phage genomic DNA isolation

The Phenol-Chloroform-Isoamyl alcohol (PCI) DNA extraction protocol (phagesdb.org) was carried out to isolate phage genomic DNA. Briefly, to remove contaminating bacterial nucleic acid from the phage preparation, a mix of 12.5 mM of MgCl_2_, 0.8 mU/ml DNAse I, and 0.01μg/μl RNase was added to phage lysate (10^10^ pfu/ml) and incubated overnight at 37 °C. After incubation, 20 mM EDTA, 50 μg/ml Proteinase K, and 0.5% SDS were added, and the mixture was incubated at 55 °C for 1 hour. An equal volume of PCI (25:24:1) was added to the lysate, mixed gently, and centrifuged at 13,000 rpm for 15 minutes, and the aqueous layer was extracted. After repeating this step three times, DNA was precipitated by adding 0.15 M sodium acetate and two volumes of 95% chilled ethanol and was kept at -20 °C. Precipitated DNA was collected by centrifugation under the same conditions, and the DNA pellet was washed with 80% ethanol, dried at room temperature, and solubilized in nuclease-free water/Tris-Cl (pH 8.0). The A260:A280 ratio and the isolated genomic DNA quantification were determined using a multimode plate reader (Biotek, India).

### 2.5 Cluster classification of isolated phages using sub-cluster-specific TMP primers

Primers (Forward: 5’-AAAGGTGATCGTGCCCATCG-3’ and Reverse: 5’-GAACCTCGTGAACAGGTCGG - 3’) were synthesized at Integrated DNA Technologies, India. Using the genomic DNA of the isolated phages as the template, cluster classification was carried out by PCR, using B1 sub-cluster-specific primers (14). PCR reactions were set up at the suggested annealing temperature (55 °C) using *Taq* DNA polymerase, and the amplicons were analyzed by agarose gel electrophoresis.

### 2.6 Determining methylation status of phage genomes

Restriction digestion of phage genomic DNA was carried out with endonucleases DpnI and DpnII (1U/μg) at 37 °C, according to the protocol described in our previous study (10). Digested products were analyzed by agarose gel electrophoresis.

### 2.7 Transmission Electron Microscopy (TEM)

For Transmission Electron Microscopy, phage preparations (10^9^ pfu/ml) were placed on a carbon-coated copper grid and stained with 2% phosphotungstic acid (PTA). Samples were observed using Jeol TEM 1400 (JEOL Ltd., Tokyo Japan) at 120 kv at SAIF (Sophisticated Analytical Instrument Facility) at CSIR-CDRI, India and the images were taken at a magnification of 30,000X.

### 2.8 Lysogenic activity of the isolated B1 mycobacteriophages

#### 2.8.1. Patch plate assay

To confirm the lysogenic status of B1 phages, a phenotypic test was performed (27). The plaques of all ten phages were obtained on *M. smegmatis* mc^2^155 after incubation at 37 °C for four days. The center of the plaques was checked for mesa formation (bacterial colonies in the centre of the plaque/spots). For each phage, a mesa colony was picked from the centre of the plaque and streaked on 7H10 plates containing Carbenicillin (50 μg/ml) and Cycloheximide (10 μg/ml) (HiMedia, India) and supplemented with 0.4% glycerol. From the streaked plate, 20 isolated colonies were patched on 7H10 plates and incubated at 37 °C for four days. Bacterial growth from each patch was resuspended in 20 μl of phage buffer and 5 μl of each suspension was spotted onto the double agar plates with top agar containing *M. smegmatis* mc^2^155 (OD_600_ of 0.4-0.6). The plates were incubated at 37 °C for 48 hours.

#### 2.8.2 Supernatant assay

From the 20 patched colonies mentioned above, one colony from each phage was inoculated in 7H9 broth, supplemented 0.05% Tween-80 and incubated at 37 °C and 200 rpm. After 48 hours of incubation, the growth obtained was used as inoculum (1%) in fresh 7H9 broth and incubated at 37 °C, 200 rpm for 48 hours again. The culture was then centrifuged at 10,000 rpm for 10 minutes, and the supernatant was sterilized using 0.2 μm filter. The filtered supernatant was spotted on 7H10 plates containing *M. smegmatis* mc^2^155 in the top agar.

### 2.9 M. smegmatis biofilm formation and treatment with phages

#### 2.9.1. Bacterial strains and growth media

For biofilm experiments, we followed a protocol (28) with minor modifications. *M. smegmatis,* strain mc^2^155 cells grown to OD_600_ 0.8 were used in 24-well plates with incubation for 96 hours at 37 °C under static condition. Quantification of biofilms was by the Crystal Violet (CV) staining method (Chakraborty et al., 2021). Briefly, after biofilm formation, the planktonic cells were removed, and the freshly prepared CV stain (1%) was added to each well. After 30-45 minutes, the stain was removed carefully, and the biofilms were washed twice with 1X PBS to remove excess stain. The stain was then extracted by 95% ethanol and the OD_595_ was measured.

#### 2.9.2. Inhibition and disruption of biofilm by B1 sub-cluster phages

To study the inhibition of biofilm formation, to wells containing *M. smegmatis* (10^6^ CFU/ml), each B1 phage (*Phages 1-10)* was added at an MOI of 100 and the plates were incubated at 37 °C for 96 hours under static conditions. For studying disruption effect of the phages, *M. smegmatis* biofilm was first allowed to form at 37 °C for 48 hours, followed by adding the phages at an MOI of 100 and extending the incubation for another 48 hours. Quantification in both the cases was done using the CV staining method.

## 3. Results and Discussion

### 3.1 Genome description of B1 cluster mycobacteriophages

B1 is one of the database’s most populated mycobacteriophage sub-cluster consisting of over 254 members out of 2257 sequenced phages. The GC content (66.5%) and codon usage profile of these phages are similar to their host *M. smegmatis.* On average, B1 phages contain about 100 protein-coding genes, share a high level of average genome identity (∼≥97%) and have a circularly permuted genome that is terminally redundant (Figure 1). The average genome length of B1 sub-cluster phages is 68,579 bp, except for phage ‘KingTut’, which has a genome length of 64,827 bp. It lacks a ∼4 kb gene fragment coding for a queuine tRNA-ribosyltransferase (QTRT), structural genes such as Head-to-tail adaptor and a few hypothetical genes in other related B1 phages. GenBank ID of phages taken as the representative in this study has been mentioned in Table S1.

**Figure 1:**
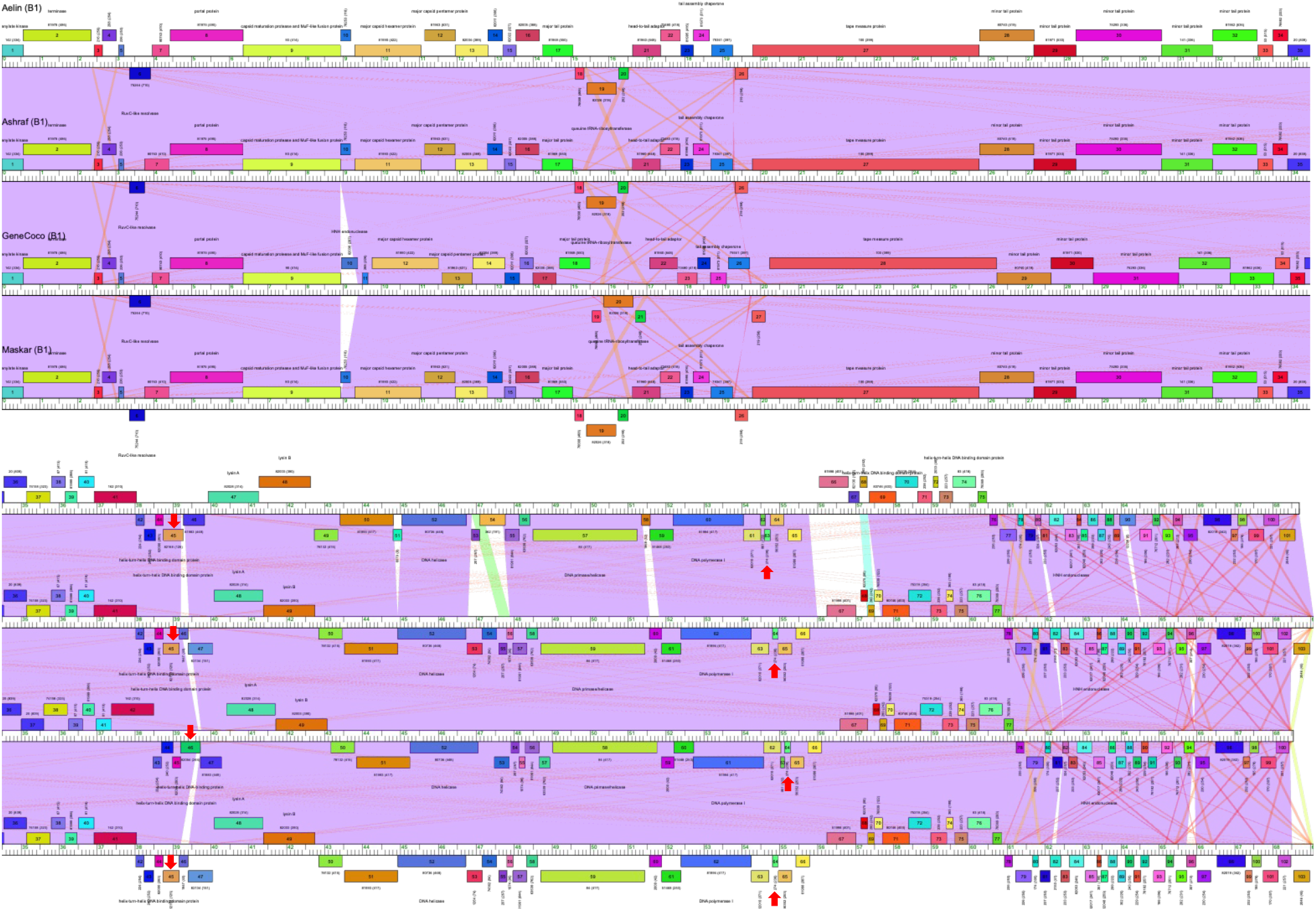
Genome comparison of representative B1 sub-cluster phages using Phamerator show a high level of sequence identity (regions depicted in purple). The average number of ORFs and genome length of the phages are approximately 100 and 68,579 bp, respectively. These phages show a unique 26 ORFs coding for hypothetical proteins at the terminal end of the reverse strand with at least ten promoter signatures. Red arrows indicate putative genes involved in lysogeny.

### 3.2 Analysis of Gene Expression Elements in B1 phage genomes

We found prediction of a high number of SigA-like promoter regions in the phage genomes of the B1 sub-cluster. At least ∼12 promoters were found in the forward strand of the genomes, and ∼29 promoters were predicted in the reverse strand. This could be attributed to an unusually high number of ORFs (50+) present in the reverse strand of the B1 genomes. Phage Aelin (MW534377) consists of 101 ORFs, of which 53 ORFs are present in the reverse strand. Similarly, in phages PDRPv & PDRPxv (10, 12), we found 57 out of 107 ORFs in the reverse strand. The reverse strand contains genes coding for RuvC-like resolvase, QTRT, the replication cassette (DNA helicase, DNA primase, DNA polymerase), DNA-binding proteins and several other hypothetical proteins. The prediction of B1 phage promoters also indicated at least three strong promoters in the region of the replication cassette, the strongest being the promoter downstream of DNA primase and upstream of the DNA polymerase (3΄-TTGACT-19 bp-CACGA). Another unique feature we observed was the presence of a continuous stretch of ∼26 ORFs coding for hypothetical proteins at the terminal end of the reverse strand (Figure 1). While the function of these proteins is not understood, it was intriguing to find at least ten top-scoring strong promoters in this cassette. All of these promoters in this stretch scored the highest among all the other promoters in the reverse strand indicating a higher transcription rate of these genes (29).

The forward strand of the genome consists of ORFs involved in genome packaging, virion structure, the endolysin cassette, and many DNA-binding and hypothetical proteins. We found approximately 12 strong promoters in this strand, such as the putative promoter 5΄-TTGACG-19 bp-GATAAC-3΄ located in the upstream region of the structural genes, which includes portal protein, capsid maturation protease, and the major capsid protein. We also predicted another promoter (5΄-TTGAGA-19 bp-CAAGAT) in the upstream region of the endolysin cassette of these phages. In an earlier study (30), we experimentally demonstrated the activity of a putative promoter region predicted upstream of the endolysin genes in PDRPxv, using the ONPG assay. Briefly, the predicted promoter region was cloned in a promoter-less vector (pSD5B) and the expression of the reporter gene (β-galactosidase) in the recombinant pSD5b confirmed the function of the putative promoter.

### 3.3 Insights into the hypothetical proteins (HPs) of B1 sub-cluster

With the growing interest in phages as antibacterial and the burgeoning sequenced phage genomes, one of the major challenges is to identify the role of a large number of ORFs, which are not understood yet. In the absence of sequence homology or motif/domain similarity, encoded products of such ORFs are deemed hypothetical proteins (HPs). However, regular attempts have been made to experimentally determine the function of these genes through the genome expression libraries (31, 32) or gene editing techniques (33, 34). Examples include the demonstration of the cytotoxic effect of hypothetical proteins (gp77-79) in L5 mycobacteriophage against *M. smegmatis* (35) and growth defects in *M. smegmatis* and *M. bovis* and *E.coli* by HPs gp34 & gp49 in mycobacteriophage D29 and Che12 (36) among others. Such efforts toward establishing the role of HPs (especially those encoding lethal genes for their bacterial host) not only insinuates the inter-species therapeutics role of phage proteins but also highlights phages as an inexhaustible and inexpensive source of antibacterial agents. Importantly, deciphering of the uncharacterized parts of phage genomes also helps us get a deeper understanding of phage biology. However, given the large number of phage genes to be decoded, relying on these molecular techniques alone can be time-consuming and labor-intensive and also requires optimization for each phage-host system. Hence, to decode the ‘dark matter’ in mycobacteriophages, we carried out a comprehensive curated computational analysis of the HPs from the B1 sub-cluster by using a combination of online computational tools with an aim to reexamine the hypothetical genes and assign putative functions.

B1 phages, on average, consist of 100 protein-coding genes, out of which approximately 76% are hypothetical. On genome analysis by DNA Master (http://cobamide2.bio.pitt.edu/computer.htm), the identified HPs were downloaded from UniProt (https://www.uniprot.org/) database and subjected to HMMER analysis (37). HMMER, a tool based on Hidden Markov models, provides information about a protein by detecting sensitive distant homologs in sequence databases. Using the tool, several of the previously uncharacterized proteins were assigned functions and were found to be either membrane or structural proteins. To annotate the HPs, we subjected each sequence to TMHMM for the identification of membrane proteins (38) and to PhANNs to identify structural proteins (19). To gain additional insights into the putative role, we identified the fold pattern of the proteins using the tool PFP-FunDSeqE (21). This enabled the prediction of a few DNA-binding and toxin proteins, which were confirmed by using the tools: DP-Bind, (analysis of DNA-binding residues) (23) and ToxDL (analysis of toxicity) (22).

This framework for annotation (Figure 2) facilitated the functional prediction of about 55% of protein-coding genes, which were hypothetical on preliminary annotation (Figure 2 and Table S2). Initially, the data was curated according to the protein families ‘phams’, which collectively represented the proteome of B1 sub-cluster phages. However, we occasionally realized a fluctuation in Pham numbers, rendering our analysis futile. To avoid confusion, we selected three phages (Aelin, Childish and GeneCoco) and their hypothetical gene products as the representative uncharacterized proteome for all the B1 sub-cluster phages in the Actinobacteriophage database (https://phagesdb.org/) (Table S2). The gene products characterized in this study were found to be common in all B1 phages. Functional prediction of these uncharacterized proteins is divided into the following categories.

**Figure 2:**
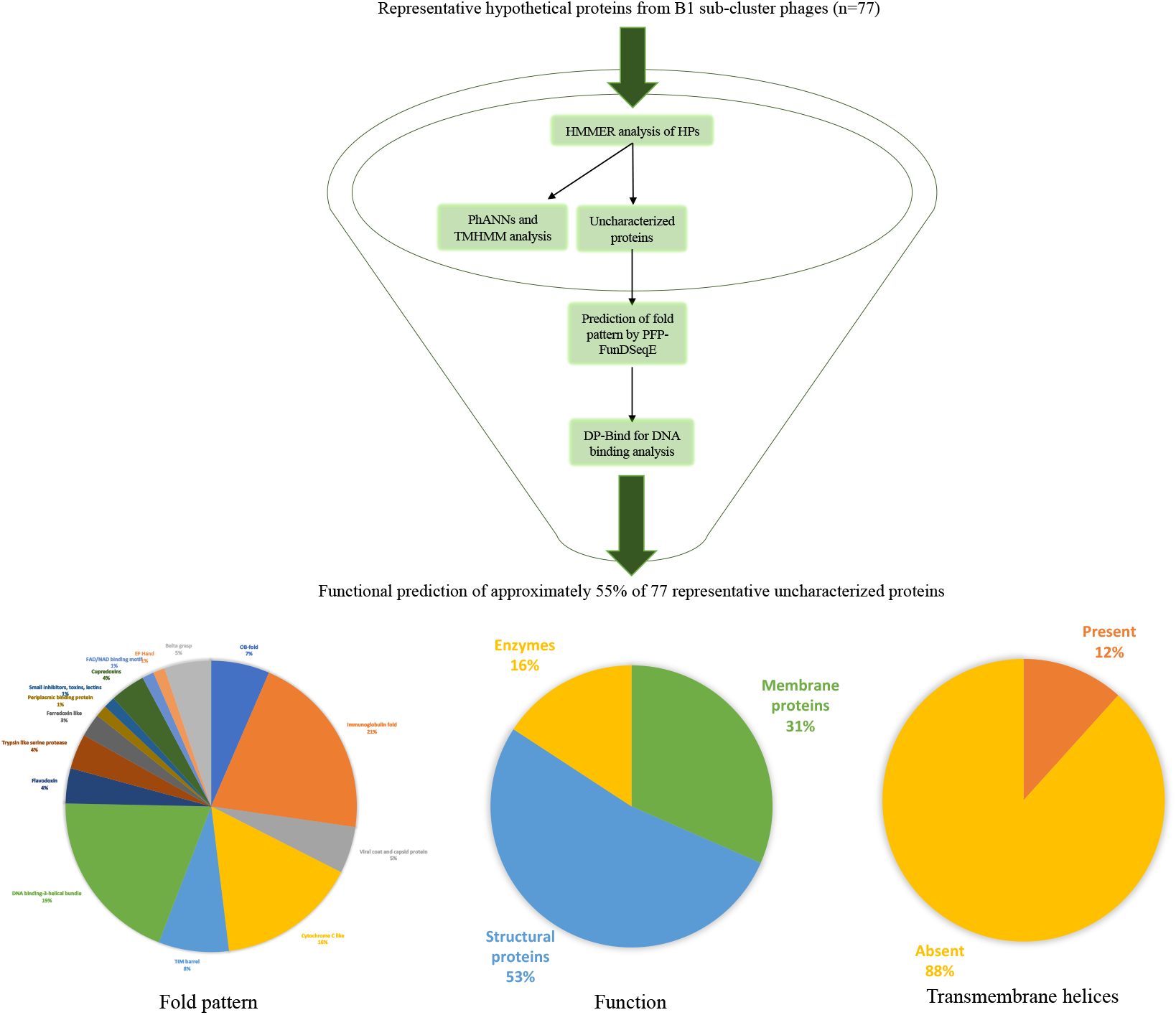
Framework of online computational tools for determination of the putative function of hypothetical proteins (HPs) in B1 sub-cluster phage genomes. This framework of annotation assisted in the functional prediction of at least 55% of additional protein ‘phamilies’, which were deemed hypothetical on preliminary annotation. Pie charts indicate the functional prediction of the HPs. Of the 77 representative HPs, 16%, 53% and 31% were predicted to be enzymes, structural and membrane proteins, respectively. Transmembrane helices were present in 12% of the HPs; immunoglobulin-like fold (21%) was most common. A detailed description of this summary has been provided in Table S2.

### 3.3.1 Membrane proteins

In all, nine putative membrane proteins were predicted (Table 1). Of these, Aelin gp7 was found to be associated with viral capsid or coat protein constituting a domain of the unknown function (DUF1360) and Aelin gp43 and Aelin gp64 were predicted to have the cupredoxin-like fold. Among the other membrane proteins, the prediction of a ‘holin-like’ protein (Aelin gp14) in B1 sub-cluster phages, which shared approximately 64% identity with the putative holin from Mycobacteriophage JAMal (sub-cluster B4) (39) appears to be unique to this cluster. The ‘holin-like’ protein (Aelin gp14) predicted here was hydrophobic with four transmembrane helices, features commonly found in phage-encoded holins (40). Although holins are an essential part of the ‘lysis cassette’ in most phages, in some, endolysins can translocate to the periplasm of the host in its absence (30, 41). The differential presence of holin-like proteins noted in B1 phages warrants further investigation.

**Table 1:** Inhibitory and disruptive effect (in percentage) of *Phages 1-10* on *M. smegmatis* mc^2^155 biofilm.

| *Phages* | **Inhibition (%)** | **Disruption (%)** |
| --- | --- | --- |
| *1* | 57.96 ± 0.038 | 29.19 ± 0.164 |
| *2* | 57.56 ± 0.063 | 46.36 ± 0.091 |
| *3* | 56.97 ± 0.141 | 19.54 ± 0.391 |
| *4* | 52.89 ± 0.041 | 25.2 ± 0.212 |
| *5* | 52.14 ± 0.041 | 45.51 ± 0.181 |
| *6* | 31 ± 0.250 | 28.15 ± 0.208 |
| *7* | 45.64 ± 0.133 | 29.36 ± 0.033 |
| *8* | 54.12 ± 0.395 | 26.03 ± 0.262 |
| *9* | 63.63 ± 0.148 | 44.33 ± 0.363 |
| *10* | 58.85 ± 0.145 | 36.79 ± 0.290 |

#### 3.3.2 Structural Proteins

Mycobacteriophages, like other bacteriophages, consist of head and tail structural proteins, forming a complete virion. The major head and minor head, along with scaffolding proteins, are the components of the head forming system, while the major tail and minor tail, along with the tape measure protein together, are a part of the tail forming system. These sets of proteins in phages are mostly present in clusters across the genome. Phages from the B1 sub-cluster also constitute the head and tail forming systems; however, some gene products in these clusters remain uncharacterized.

HMMER and PhANNs predicted total ten structural proteins (Table S1). We re-annotated and discovered ORFs coding for a minor tail protein (Aelin gp13, gp34, gp36, gp37 and gp41), major tail protein (gp25 and gp40), head-to-tail connector (gp18), tail fibre (gp33) and tail assembly chaperone (gp23). However, the re-annotation of gp25 resulted in contradictory outcomes. HMMER predicted the ORF to code for a tail assembly chaperone (3.5e-54), while PhANNs predicted the protein to be a major tail with 57% confidence. In such ambiguous cases of structural proteins, their purification and biophysical characterization can help ascertain their roles.

#### 3.3.3 Proteases and Nucleases

Phage-encoded enzymes perform a myriad of essential functions in their life cycle. Enzymes such as endolysins assist in the lysis of the host cell wall, while proteases and nucleases perform the maturation of the head-forming unit and efficient packaging of the genome, respectively. Phage-encoded proteases have been exhibited to play a key role in the processing of the major capsid. The use of HMMER and PFP-FunDSeqE assisted in the prediction of nucleases and proteases among the hypothetical proteins of these phages. Aelin gp37 and gp38 were found to have trypsin-like serine protease folds. Enzymes with this fold have been demonstrated to cleave the polypeptide chains by acting on the positively charged (Arg/Lys) amino acid residues (42). This enzyme assists in the conformational change of the procapsid precursor to give rise to a mature or major capsid protein of the bacteriophage (43). Therefore, Aelin gp37 and gp38 with a serine protease fold could play a similar role in B1 phages.

Phages encode several nucleases that are responsible for functions such as cleavage of cytosine-containing host DNA, correct packaging of viral DNA in the capsid, and inducing site (non)specific recombination (44–46). While re-annotating the HPs, we predicted the presence of HNH endonucleases (Aelin gp54) consisting of the characteristic histidine and asparagine residues. These enzymes are responsible for homologous recombination in phages by nicking the double-stranded DNA, which causes gene conversion (47). The enzymes with HNH motif can either have non-specific nuclease activity or may consist of conserved DNA recognition domains. They could play a role in the observed mosaicism in mycobacteriophage genomes (48).

#### 3.3.4 Other proteins

We could also assign a function to a few more uncharacterized proteins. PFP-FunDSeqE tool predicted several HPs (Aelin gp20, gp53, gp62, gp67, gp70, gp75, gp76, gp80, gp84, gp86, gp88, gp91, gp93, gp94 and gp95) to have a fold pattern similar to DNA binding-3-helical bundle (Table S1). To confirm this annotation, we analyzed the sequences by the DP-Bind tool, which confirmed the presence of DNA binding residues. Proteins with this folding pattern, consisting of an HTH DNA-binding domain, play significant roles, such as regulation of transcription and lytic/lysogenic growth, replication, and DNA repair in prokaryotes (49). Annotation of the DNA binding-3-helical bundle fold pattern and confirmation of DNA binding residues can therefore indicate similar functions of these uncharacterized proteins. Besides the DNA-binding proteins, HMMER annotation predicted Aelin gp5 to be a zinc finger protein. Such proteins have been demonstrated to bind to mRNA and act as a translational factor in bacteriophages (50). Another gene product Aelin gp72 was annotated as a CHAT domain-containing protein (9.3e-06). CHAT for the association of Caspase HetF with tetratricopeptide repeat (TPR) motif-containing proteins. Proteins of this family are related to peptidases, which include caspases (IPR024983). Other than this, we also found several HPs to have an immunoglobulin and cytochrome c-like fold (Table S1).

### 3.4 Insights into genetic evolution of B1 sub-cluster phages

Mycobacteriophages from cluster O have been previously observed to show signs of evolution due to deletions and insertions (indels) in their genome (51). On analyzing B1 sub-cluster phages, the observed difference in genome compositions could be due to horizontal gene transfers or indels, suggesting their evolution from a common ancestor.

#### 3.4.1 Evidence of horizontal gene transfer

The horizontal transfer of genes between the host bacteria and bacteriophages has been reported to be one of the reasons behind the genetic evolution and mosaicism of phage genomes (51). However, gene transfer from eukaryotic organisms is a unique way of acquisition of genetic material in prokaryotes which could also indicate plausible genetic evolution in phages. This method of gene transfer has been reported earlier in phages infecting *Listeria* and *Wolbachia* (52, 53). Through HMMER analysis, we found two such gene products in B1 sub-cluster phages that potentially indicate horizontal gene acquisition from eukaryotes. Aelin gp20 was similar to an Ankyrin Repeat Region domain-containing protein (1.7e-06) predominantly found in eukaryotes. They can bind to other proteins and are involved in diverse functions such as ion transport, protein kinase activity, etc. (54). Another protein, Aelin gp71, is a structural motif annotated as a Tetratricopeptide (TPR) repeat (5.7e-05). TPRs are responsible for many protein–protein interactions in eukaryotic processes, such as apoptosis, cell signaling and inflammatory response. Also, effector proteins of certain intracellular pathogens of eukaryotes frequently contain TPR repeats and therefore hint at a eukaryotic origin. We don’t know the role of these genes, phages harbouring them suggests they might have infected the intracellular pathogenic bacteria of eukaryotic cells in the past (53).

#### 3.4.2 Evidence of Insertions and Deletions

Even though a high identity is observed in the genome sequence of sub-cluster B1 phages, the selective presence of specific ORFs in phage genomes can be taken as evidence for insertion and deletion (indel) events. For example, Gp10 in phage Chorkpop, which shows 94% identity with the HNH endonuclease of phage OliverWalter, is an insertion, absent in several other genomes (Figure 1). Likewise, Chaelin Gp26, a unique stretch of 105 bp, appears to belong to an entirely new family with no similarity with any other phage protein and Gp46 in Andre phage codes for an uncharacterized protein, which shows no significant similarity with any other proteins in the database. Examples for deletions include Gp46 in Ashraf, Maskar and Andre is a hypothetical membrane protein absent in the genomes of Aelin, Chaelin, Bishoperium, Chorkpop, etc. It will require a larger-scale analysis of phage genomes from different clusters to understand the significance of the indels in phage evolution, beyond their role in creating genome sequence diversity.

### 3.5 Isolation of B1 mycobacteriophages from soil samples

The method for phage isolation was the soft agar overlay technique, where the filtered enriched soil lysate was plated on the *M. smegmatis* (mc^2^155) lawn. A total of 160 soil samples were screened, 30 mycobacteriophages were isolated and purified and studied. High-titre (10^10^-10^12^ pfu/ml) of each phage was prepared, aliquoted, stored and processed for further analysis.

### 3.6 B1 sub-cluster phages

A study by Smith et al. 2013 put forth an elegant single-gene analysis method that can establish cluster relationships in phages. They used conserved regions within the tape measure protein (TMP) gene to classify phages into clusters and sub-clusters. We used B1 sub-cluster-specific unique primers to identify B1 sub-cluster phages. Based on the size of the amplicon (493 bp), ten phages were found to belong to the B1 sub-cluster (Figure 3). Henceforth, these B1 phages are denoted as *Phages 1-10*.

**Figure 3:**
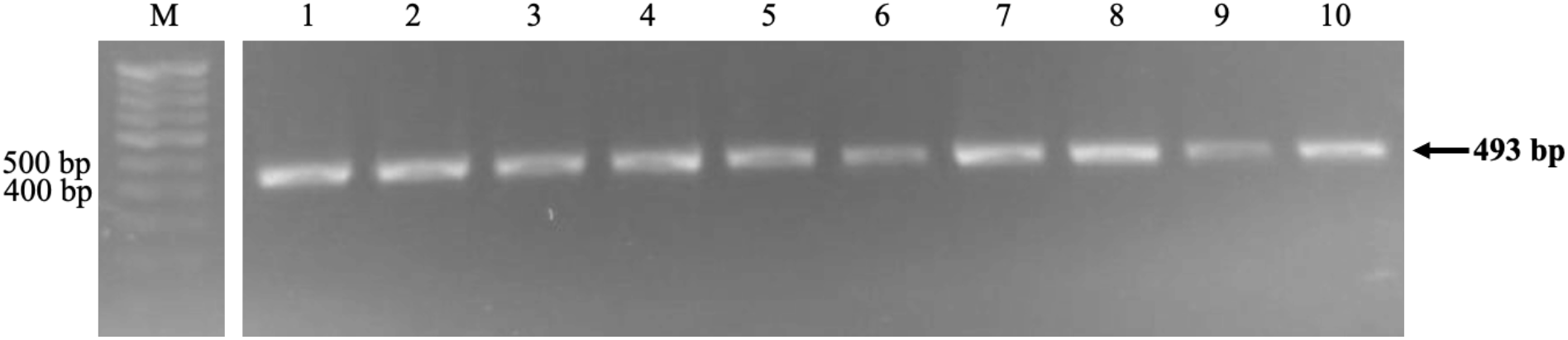
PCR analysis using B1 sub-cluster specific TMP gene primers. M: Marker 100 bp, Lane 1-10: *Phages 1-10*. An amplicon size of 493 bp was observed in all the ten phages confirming them to belong to the B1 sub-cluster.

#### 3.7 Methylation status of the genomes

The bacterial host’s restriction-modification system (RMS) is a key cellular defense mechanism against phage infection. A methylated (modified) phage genome hints phage either carries its methylase or survived restriction digestion in the previous host by chance modification of its DNA along with that of the host. On restriction digestion with DpnI, which cleaves at the methylated restriction site and DpnII, which cannot cut at the methylated sites, we found that all ten B1 mycobacteriophage genomes are non-methylated (Figure 4). We also did not find a gene encoding cytosine-C5-specific DNA methylase in B1 phage genome. Taken together, B1 sub-cluster phages appear to possess a different defense mechanism.

**Figure 4:**
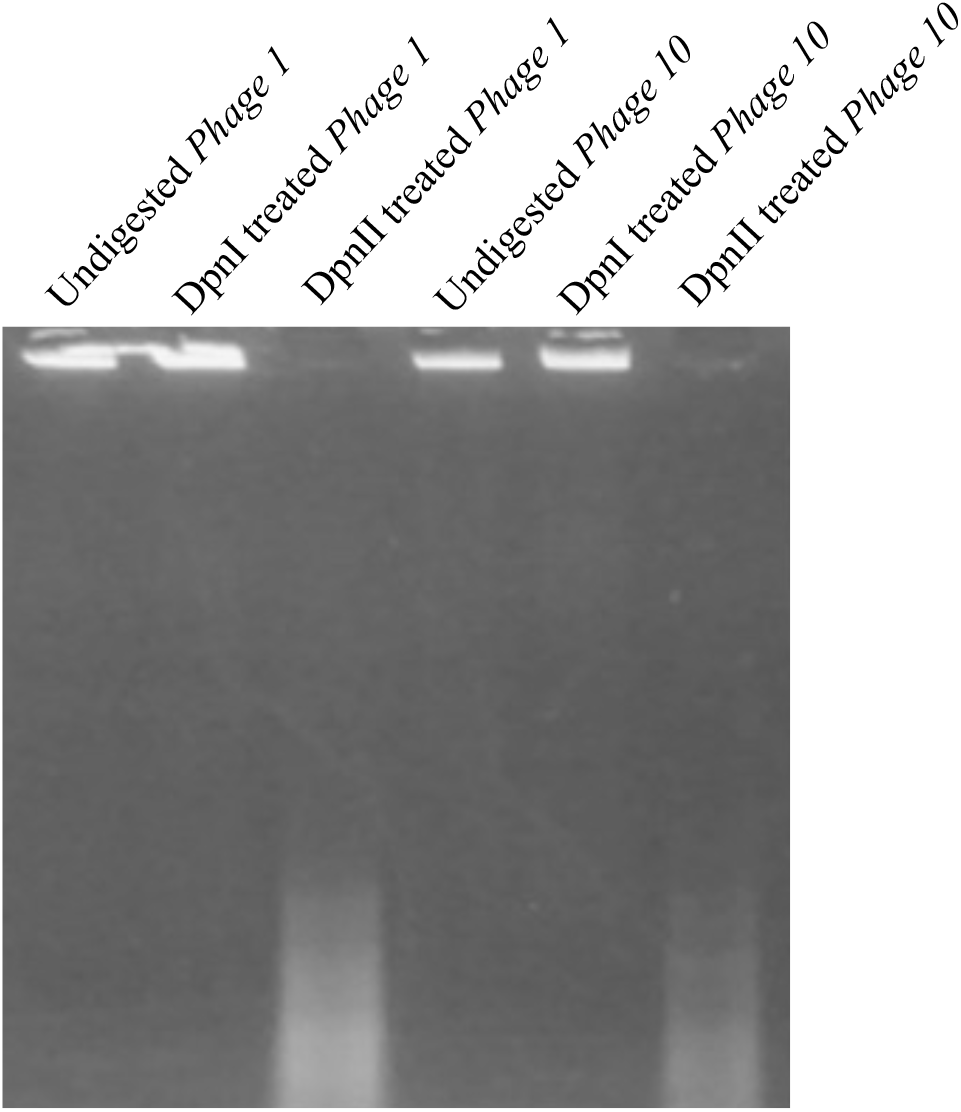
Agarose gel electrophoresis of two B1 phage genomes treated with DpnI, which can cleave at the methylated restriction site and DpnII enzyme, which cannot cut at the methylated sites. Lane 1 and 4: Uncut genomic DNA of *Phage 1* and *Phage 10*. Lane 2 and 5: Genomic DNA of *Phage 1* and *Phage 10* treated with DpnI. Lane 3 and 6: Genomic DNA of *Phages 1* and *10* treated with DpnII. Data not shown for the other 8 phages.

#### 3.8 Morphology of B1 sub-cluster phages and their plaques

The ten B1 phages showed varied plaque morphology, which included circular and bull’s eye and clear or turbid plaques (Figure 5A). Plaque sizes ranged from 0.53 cm to 0.25 cm in diameter (Figure 5B). Halo formation (0.2 - 0.3 cm) in all the phages was observed after 72 hours of incubation. A halo around the plaque is usually attributed to an exopolysaccharide (EPS)-degrading enzyme, which can also aid in the antibiofilm activity of the phages (55).

**Figure 5:**
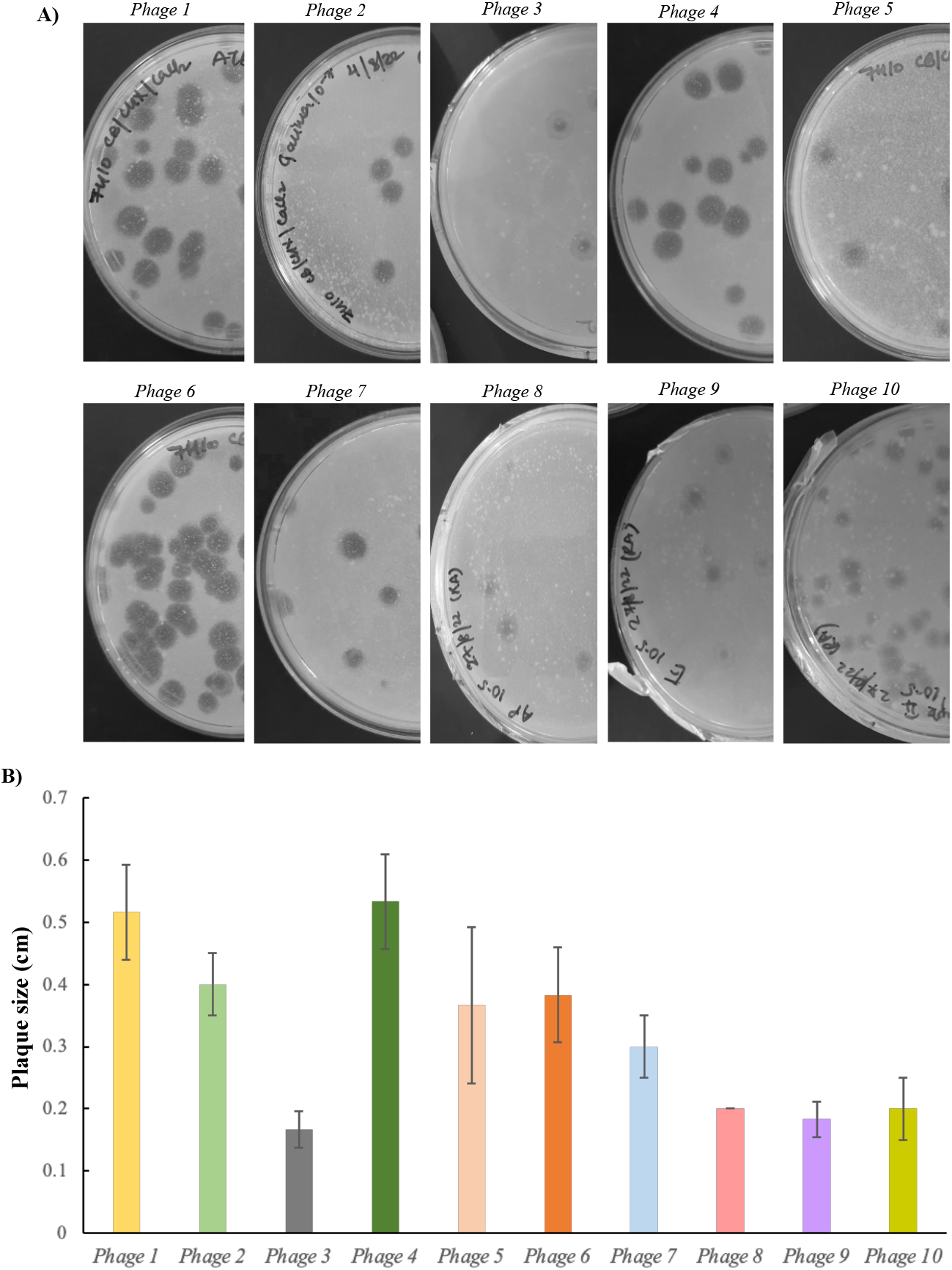
Plaque morphology of B1 phages. The size of five plaques from each phage was measured, and the average is plotted here. *Phages 1-10* showed different plaque morphology and size upon incubation. The largest and smallest plaque size is observed in the case of *Phage 4* (0.7 ± 0.1 cm) and *Phage 3* (0.1 ± 0.02 cm), respectively. Halo formation in all phages was observed after 72 hours of incubation, and the size of the halo varied in all phages (0.2 - 0.3 cm).

Transmission electron microscopy showed the virions to have icosahedral heads and long non-contractile tails (Figure 6A). The average head and tail diameter were 51.33 nm and 202.12 nm, respectively (Figure 6B).

**Figure 6:**
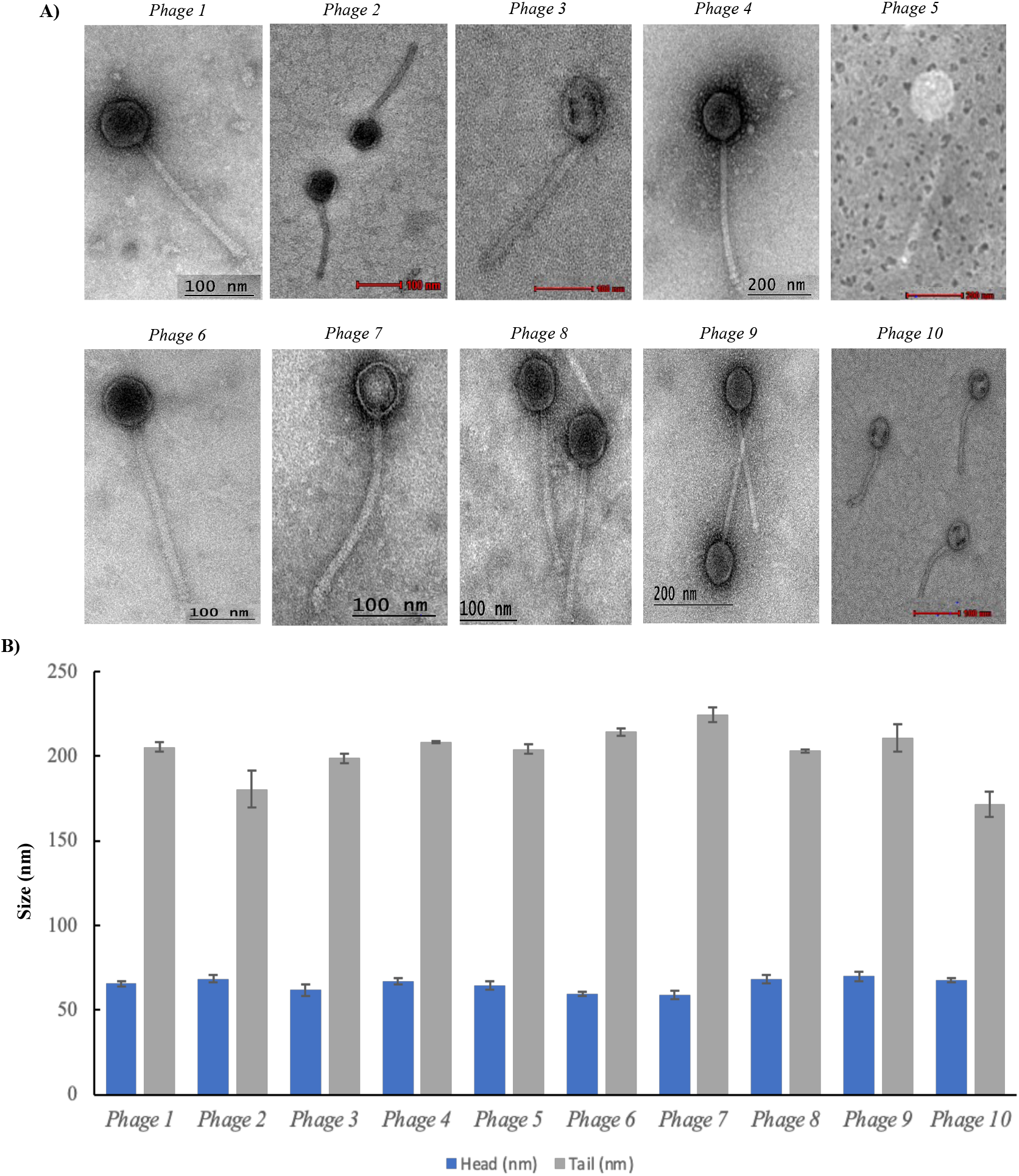
Virion morphology and size of the characterized B1 mycobacteriophages. **A)** Electron micrograph of the isolated B1 phages shows an icosahedral head and a long non-contractile tail. **B)** Head and tail length of B1 sub-cluster phages (in nm) at 100 or at 200 nm scale. TEM analysis of purified phages showed an average head and tail size of 65 and 202.12 nm, respectively.

### 3.9 B1 sub-cluster phages show unstable lysogenic behavior

The presence of repressor, integrase and excisionase genes in the phage genome (56) typically suggests the phage to lysogenize the bacterial host. In mycobacteriophages, the lysogeny has been observed in A, F, Q and N clusters (57), but in B cluster phages, stable lysogens have not been reported (1). In our previous reports on the B1 sub-cluster phage ‘PDRPxv’ we predicted the presence of a repressor protein (gp47) and a Recombination Directionality Factor (RDF) or Xis protein (gp68), which are known to be involved in lysogeny (10). Gp47 belong to the xenobiotic response element (XRE) family of transcription regulators which has been experimentally demonstrated to act as a repressor protein in phage L5 (58) and RDF protein (gp68) or Xis, directs the excision of prophage from the host genome, at attL and attR sites (59). In this study, we have also predicted these proteins to be encoded by other B1 sub-cluster phages (Figure 1). Although these phages lack an integrase, the presence and the precise role of other genes responsible for lysogeny could not be discerned. Therefore, we examined the genomes for other modes of lysogeny (if any) and, on comparative genome analysis, found a putative DNA primase/helicase, showing sequence similarity to RepA-like helicases encoded by phages from B5 and B13 sub-cluster (Figure 7A). This was also supported by the results from domain analysis, where a hexameric replicative helicase RepA domain was predicted in the proteins mentioned above (Figure 7B). Temperate phages that lack an integrase but code for RepA-like proteins have been previously demonstrated to be necessary and sufficient for extrachromosomal replication of prophage DNA within their host (60). From this analysis, we speculate the phages from the B1 sub-cluster to undergo a RepA-mediated lysogeny cycle wherein the prophage DNA is replicated extra chromosomally without the need for an integrase. To confirm it experimentally, we performed an assay developed by Altamirano & Barr (27) and found *Phages 1-10* to show spontaneous (without induction) release of the prophages from the *M. smegmatis* lysogens (Figure 8). This also has been observed previously for other mycobacterium prophages (61, 62). While the plasmids carrying a *repA* cassette has been previously shown to be unstable and poorly maintained in *Mycobacterium* spp. (60), our experiment analyses infers temperate life cycle of the B1 phages under study here. We also observed variations in the appearance of clear zones (or no zone at all in *Phages 2* and *5*) among the twenty lysogens (isolated from mesa from each of the ten phages) (Figure 8). This could be either due to the differential expression of the depolymerase gene or due to the varied regulatory control of phages on the gene expression of the lysogenized host (63).

**Figure 7:**
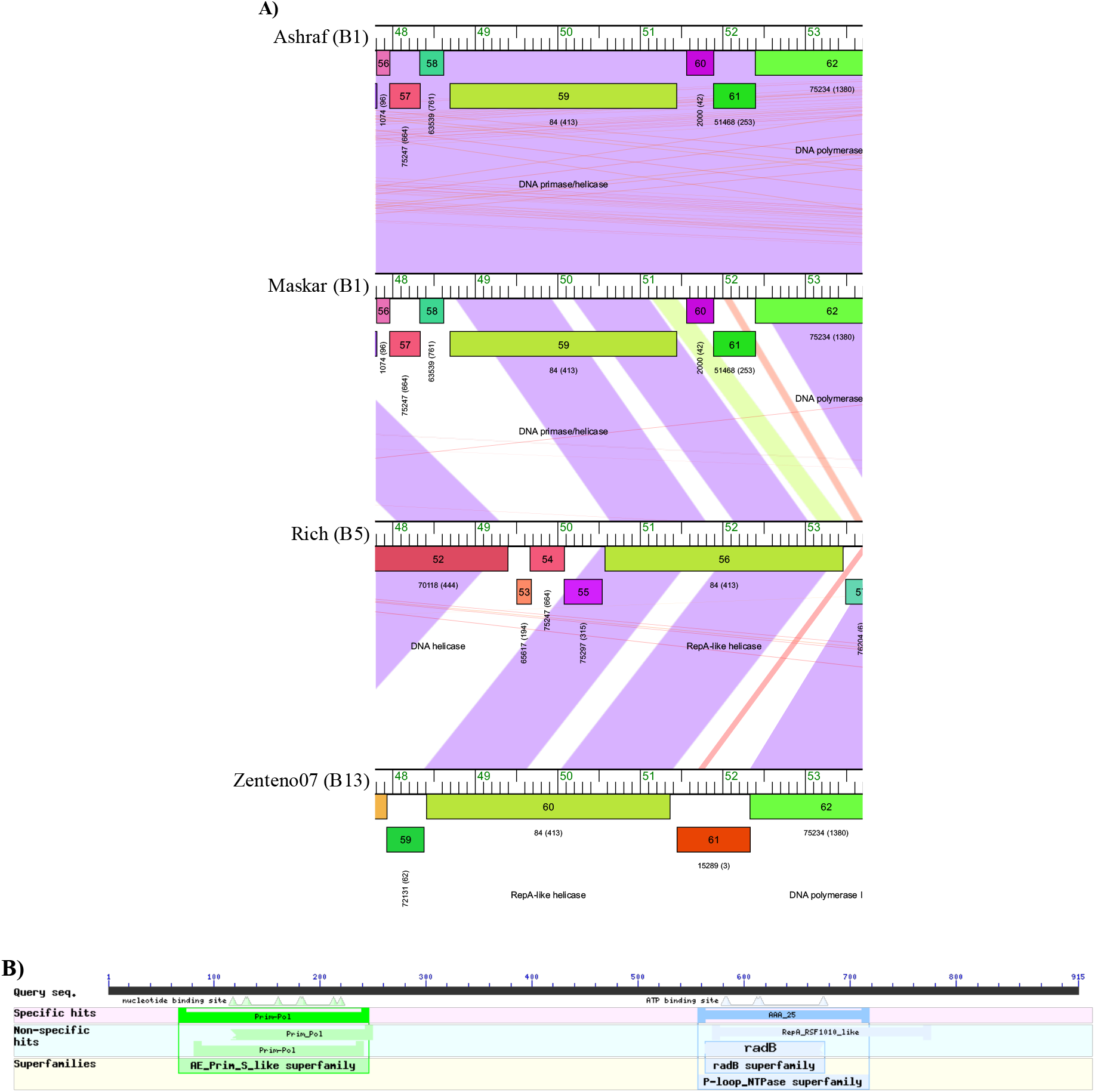
Genome comparison using Phamerator and domain analysis of representative B cluster phages. **A)** Genome comparison of B1 sub-cluster phages Ashraf and Maskar with B5 cluster phage Rich and B13 cluster phage Zenteno07 show sequence similarity of RepA-like helicase, which has been previously reported to be involved in extrachromosomal replication of phage DNA during the lysogenic cycle. **B)** Domain analysis of DNA primase/helicase from representative B1 sub-cluster phage Ashraf shows the presence of hexameric replicative helicase RepA domain (571 aa–776 aa).

**Figure 8:**
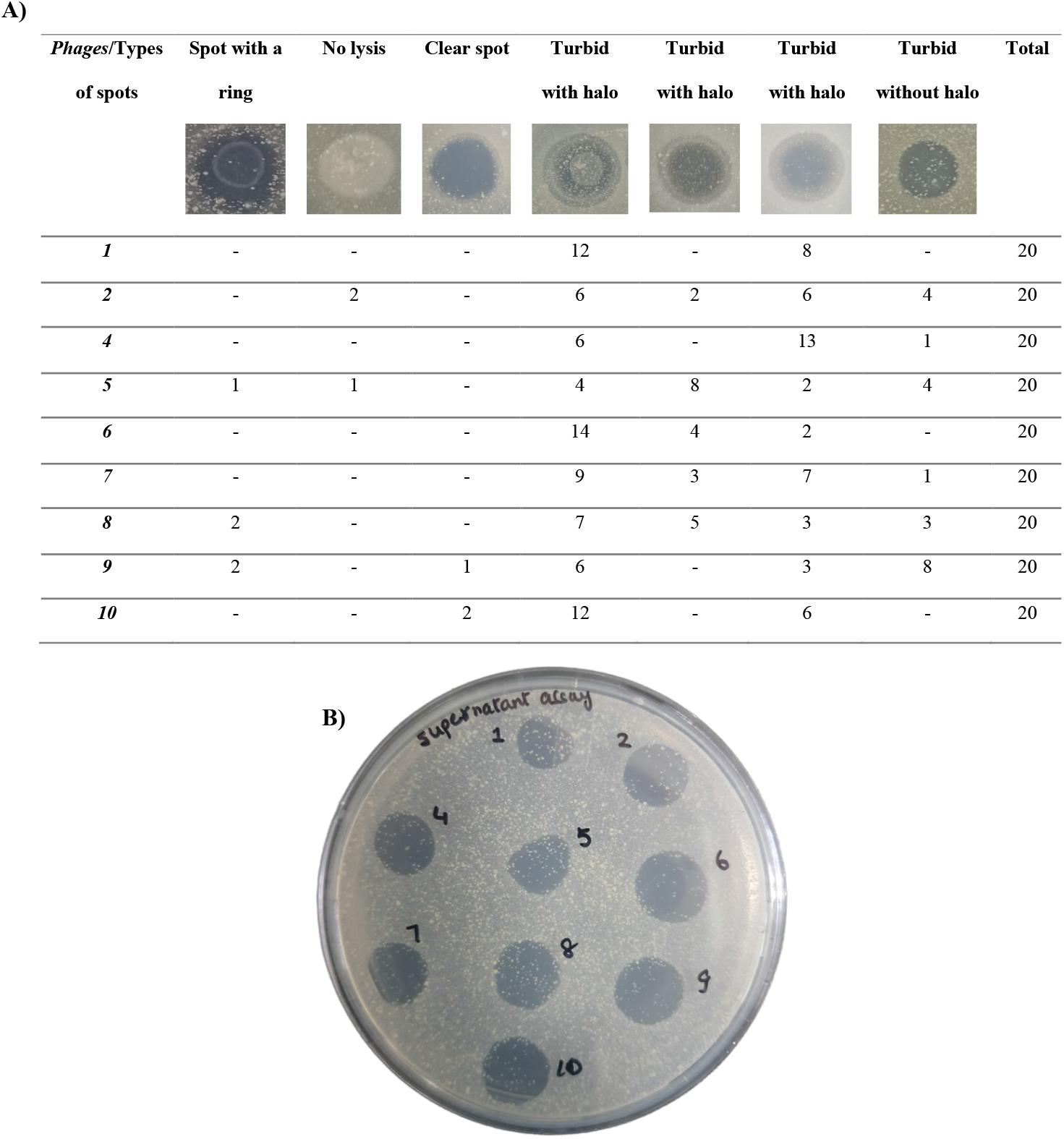
Screening for lysogenic behavior of *Phages 1-10*. **A)** Patch plate assay of the *M. smegmatis* mc^2^155 lysogens isolated from mesa colonies from each of the ten phages. The numbers given in the table depict how many times a particular spot morphology was observed out of the 20 colonies isolated from mesa from each of the ten phages. **B)** Supernatant Assay: A single lysogenized colony out of the 20 mentioned above from *Phages 1-10* was picked and grown in 7H9 media for 48 hours. The filtered supernatant from the culture was spotted on *M. smegmatis* lawn and showed the spontaneous release of phages. Both the sets of experiments were performed in duplicates.

### 3.10 B1 sub-cluster phages against M. smegmatis biofilm

#### 3.10.1 Biofilm Inhibition and Disruption by B1 sub-cluster phages

Mycobacteria can form biofilms even in the harshest conditions, adding up to their virulence and ability to act as reservoirs for various opportunistic infections (7). Infection recurrence, immune system evasion and inability to respond to antibiotic treatment are all characteristics of the disease-causing mycobacteria and biofilm formation to colonize host cells (7). However, only limited reports are available on the antibiofilm activity of mycobacteriophages (28, 64). Since phage-encoded depolymerase enzymes are reported to target extracellular or surface polysaccharides, i.e., bacterial capsules, slime layers, biofilm matrix, or lipopolysaccharides (65), we first analyzed the structural proteins of B1 phages using PHYPred to predict the presence of any encoded enzymes that could aid impacting the biofilm. Minor tail protein (Aelin, Maskar, Ashraf gp32) and a hypothetical protein (Aelin, Maskar, Ashraf gp40) were predicted as enzymes among the other structural proteins with probability scores of 0.56 and 0.52, respectively. We used the same tool to analyze if the predicted enzymes were hydrolase. Gp40 was predicted to be a hydrolase, and gp32 was not. Since phage hydrolases can be ectolysin and not depolymerase (65), gp32 was further examined using the I-TASSER tool, which predicted the protein as a pectin depolymerase (GO score: 0.6, GO:0004650) with a biological function in carbohydrate metabolism (GO score: 0.6, GO:0005975).

Based on the predicted presence of a depolymerase-like enzyme in B1 phages, we tested the antibiofilm activity of *Phages 1-10 in vitro*. Liquid-air interface biofilms of *M. smegmatis* mc^2^155 were formed in 24-well plates, and phages were tested for their ability to inhibit and disrupt biofilm (Figure 9A-C). The average OD_590_ of untreated *M. smegmatis* biofilm (after the CV assay) was 1.6-2.6 (Figure 9D). *Phages 1-10* showed varying levels of inhibition and disruption of the host biofilm (Table 1) with the highest inhibition (64%) demonstrated by *Phage 9* (Figure 9D), and the highest disruption (46%) was caused by *Phage 2* (Figure 9E). Though the observed antibiofilm effect of B1 Phages could not be solely attributed to putative occurrence of depolymerases, nevertheless, it is a promising data to get.

**Figure 9:**
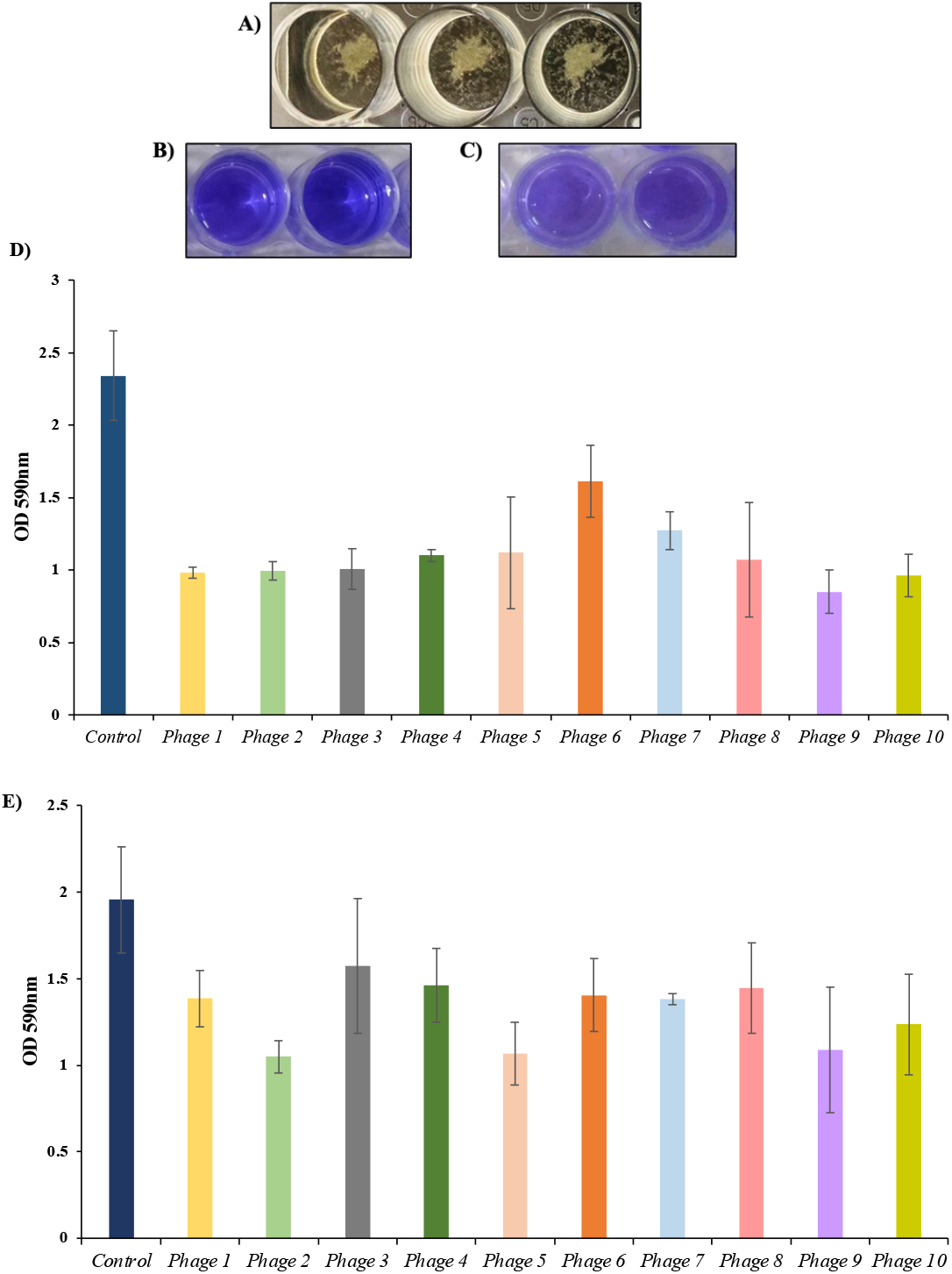
Inhibition and disruption activity of B1 sub-cluster mycobacteriophages (*Phages 1-10*) *M. smegmatis* mc^2^155 biofilm. **A)** *M. smegmatis* liquid-air interface biofilm formation. **B)** *M. smegmatis* biofilm (control, without the addition of phages), after 96 hours of incubation at 37 ℃ and staining with crystal violet. **C)** An example of *M. smegmatis* biofilm, treated with phage at an MOI of 100 after 96 hours of incubation at 37 ℃ and stained with crystal violet. **D)** Inhibitory activity of *Phages 1-10* on *M. smegmatis* biofilm after 96 hours of incubation at 37 ℃. Control-*M. smegmatis* mc^2^155 cells; Treatment groups: *Phages 1-10*. **E)** Disruption activity of *Phages 1-10* (10^8^ PFUs/ml) on *M. smegmatis* mc^2^155 biofilm after 96 hours of incubation at 37 ℃. Control-*M. smegmatis* mc^2^155 cells; Treatment groups: *Phages 1-10.* Each experiment was performed twice in triplicates, and the result was calculated as the mean percentage of the inhibition observed.

### 3.11 Phages are effective against drug-resistant M. smegmatis

The lipid-rich, waxy cell wall of mycobacteria spp is hard to penetrate, rendering them naturally immune to a wide range of drugs and also linked to opportunistic infections in humans and animals (66). *M. smegmatis,* although are considered non-pathogenic, they can cause opportunistic infections. Wallace *et al.* and colleagues reported the presence of drug resistance in 22 human isolates of *M. smegmatis*, out of which 19 were from skin or soft-tissue infections. These isolates were resistant to isoniazid (INZ) and rifampin, the first line of anti-tubercular drugs, which in some cases posed a severe threat (67). Recently, there has been a rise in INZ-resistant mycobacterial infections globally, which can impact the effectiveness of INZ-based therapy for mycobacterial infections. In a recent study (68), the efficacy of mycobacteriophages against INZ-resistant *M. smegmatis* was demonstrated and, PDRPv and PDRPxv (B1 sub-cluster phages) were shown to reduce bacterial load significantly (2).

Although none of the ten B1 phages showed activity against *M. tuberculosis* H37Rv and *M. fortuitum* (NTM) in our study, they were found to kill the isoniazid (INZ)-resistant strain of *M. smegmatis* (4XR1). Phage activity against 4XR1 was observed in spot test as well as in plaque assay (data not shown), which hints toward their application for containing opportunistic infections caused by drug-resistant *M. smegmatis* isolates.

## Conclusion

We have characterized B1 sub-cluster mycobacteriophages, revealing a range of new insights at genomic as well as phenotype levels. Even though the number of sequenced mycobacteriophage genomes has increased exponentially over the last decade, information regarding their characteristic traits comes from very few. Here, we explore the genomic cues of the B1 sub-cluster phages in depth, by analyzing the genome sequences already available in the database and also show the morphological and life cycle features and application of B1 phages as anti-*M. smegmatis* biofilm agents, experimentally. We believe an adequate knowledge on phages for other bacterial phages too can be achieved when computational and experimental analysis is conducted in conjunction, as shown in this study.

## Acknowledgement

We thank the Indian Council of Medical Research (ICMR), New Delhi, India, for supporting the project (5/8/5/38/2019-ECD-I). RA received the Senior Research Fellowship (SRF) from the University Grants Commission (UGC), New Delhi, India; KN received the Innovation in Science Pursuit for Inspired Research (INSPIRE) fellowship from the Department of Science and Technology (DST), India. SS received the Project Assistant Fellowship from another project. SKR received fellowship from Council for Scientific and Industrial Research (CSIR), India. Acharya Narendra Dev College (ANDC), University of Delhi, is gratefully acknowledged for providing the infrastructural facilities and ELITE fellowship to RD.

## Author contributions

UB conceptualized, designed, and supervised the experiments and edited the manuscript. RD, RA, KN and SS performed the experiments and wrote the manuscript. RD performed the computational analysis. RA and KN contributed equally and are both second authors. AKS and SAP performed phage screening on *Mycobacterium tuberculosis* (H37Rv). SKR and AM performed the TEM analysis of phages. All authors read and approved the final manuscript.

## Availability of data and materials

All data generated or analyzed during this study are included in this published article and its supplementary information files.

## Consent for publication

Not applicable.

