## Supplemental Table 1 and 2 for "Insights into the genomic features, lifestyle and therapeutic potential of B1 sub-cluster mycobacteriophages"

**Address for communication:**

Urmi Bajpai, PhD (Professor)

Department of Biomedical Science,

Acharya Narendra Dev College (University of Delhi),

Govindpuri, Kalkaji, New-Delhi-110019, India.

**Supplementary Information**

**Table S1:** List of B1 phages from database examined in the genome analysis study and their GenBank accession numbers.

| Phage designation | GenBank Accession |
|---|---|
| KingTut | MH450122 |
| PDRPv | KR029086.1 |
| PDRPxv | KR029087.1 |
| L5 | Z18946 |
| D29 | AF022214 |
| Che12 | DQ398043 |
| Chorkpop | KY676783 |
| OliverWalter | MG925356 |
| Chaelin | MT310867 |
| Ashraf | KY385380 |
| Maskar | KY385383 |
| Aelin | MW534377 |
| Bishoperium | MK279897 |
| Childish | MH371114 |
| GeneCoco | MH479916 |

**Table S2:** Functional prediction of hypothetical proteins (HPs) from B1 sub-cluster using the framework for annotation proposed in this study. Gene products from phages Aelin (MW534377), Childish (MH371114) and GeneCoco (MH479916) are used as representatives for all B1 sub-cluster phages.

| Gene product (gp) | HMMER Prediction (e-value) | PhANNS Prediction (Confidence) | Transmembrane Helices | Fold Pattern | DNA Binding Residues | Toxicity |
|---|---|---|---|---|---|---|
| **Aelin gp3** | - | - | - | OB-fold | N/A | Non-toxic |
| **Aelin gp4** | - | - | - | Immunoglobulin-like | N/A | Non-toxic |
| **Aelin gp5** | Zinc finger protein OZF-like (3.8e-07) | - | - | Immunoglobulin-like | N/A | Non-toxic |
| **Aelin gp7** | Membrane protein (DUF1360) (1.4e-26) | - | Present | Viral coat and capsid protein | N/A | Non-toxic |
| **Aelin gp10** | - | - | - | Cytochrome C like | N/A | Non-toxic |
| **Aelin gp12** | - | - | - | Cytochrome C like | N/A | Non-toxic |
| **Aelin gp13** | - | Minor tail protein (72%) | - | Cytochrome C like | N/A | N/A |
| **Aelin gp14** | Holin (1.5e-35) | - | Present | OB-fold | N/A | Non-toxic |
| **Aelin gp15** | Membrane Protein (1.7e-37) | - | Present | Cytochrome C like | N/A | Non-toxic |
| **Aelin gp16** | Membrane Protein (2.2e-57) | - | Present | TIM barrel | N/A | Non-toxic |

| | | | | | | |
|---|---|---|---|---|---|---|
| **Aelin gp18** | - | Head-tail joining protein (68%) | - | Cytochrome C like | N/A | N/A |
| **Aelin gp20** | Ankyrin Repeat Region domain (1.7e-06) | - | - | DNA binding-3-helical bundle | Present | Non-toxic |
| **Aelin gp22** | - | - | - | Immunoglobulin-like | N/A | Non-toxic |
| **Aelin gp23** | Tail assembly chaperone (9.3e-11) | - | - | Immunoglobulin-like | N/A | N/A |
| **Aelin gp25** | Tail assembly chaperone (3.5e-54) | Major tail (57%) | - | Flavodoxin-like | N/A | N/A |
| **Aelin gp26** | Putative outer membrane protein (1.2e-05) | - | - | OB-fold | N/A | Non-toxic |
| **Aelin gp33** | - | Tail fibre (65%) | - | Trypsin-like serine protease | N/A | N/A |
| **Aelin gp34** | Minor Tail protein (2.0e-07) | - | - | EF-hand | N/A | N/A |
| **Aelin gp36** | Minor Tail protein (4.8e-66) | Minor tail protein (89%) | - | Immunoglobulin-like | N/A | N/A |
| **Aelin gp37** | Minor Tail protein (9.5e-138) | - | - | Trypsin-like serine protease | N/A | N/A |
| **Aelin gp38** | - | - | - | Viral coat and capsid protein | N/A | N/A |

| | | | | | | |
|---|---|---|---|---|---|---|
| **Aelin gp39** | - | - | - | Ferredoxin-like | N/A | |
| **Aelin gp40** | Structural Protein (3.2e-36) | Major tail (58%) | - | TIM barrel | N/A | N/A |
| **Aelin gp41** | Structural Protein (3.9e-127) | Minor tail protein (67%) | - | Cytochrome C like | N/A | N/A |
| **Aelin gp42** | - | - | - | Immunoglobulin-like | N/A | N/A |
| **Aelin gp43** | - | - | Present | Cupredoxins | N/A | N/A |
| **Aelin gp46** | HTH Domain (2.0e-42**)** | - | - | Flavodoxin-like | Present | N/A |
| **Aelin gp49** | - | - | - | Periplasmicbinding protein | N/A | N/A |
| **Aelin gp50** | PD-(D/E)XK-1 Domain (9.6e-136) | - | - | Immunoglobulin-like | N/A | Non-toxic |
| **Aelin gp51** | - | - | - | Immunoglobulin-like | N/A | Non-toxic |
| **Aelin gp53** | - | - | - | DNA binding-3-helical bundle | Present | Non-toxic |
| **Aelin gp54** | HNH endonuclease (9.4e-131) | - | - | Immunoglobulin-like | N/A | N/A |
| **Aelin gp55** | Membrane Protein (2.8e-19) | - | Present | Immunoglobulin-like | N/A | Non-toxic |
| **Aelin gp56** | - | - | - | Cytochrome C like | N/A | Non-toxic |
| **Aelin gp58** | - | - | - | Trypsin-like serine protease | N/A | N/A |
| **Aelin gp59** | - | - | - | Ferredoxin-like | N/A | Non-toxic |

| | | | | | | |
|---|---|---|---|---|---|---|
| **Aelin gp61** | - | - | - | TIM barrel | N/A | Non-toxic |
| **Aelin gp62** | - | - | - | DNA binding-3-helical bundle | Present | Non-toxic |
| **Aelin gp63** | - | - | - | Small inhibitors, toxins lectins | N/A | Non-toxic |
| **Aelin gp64** | Membrane Protein (8.7e-08) | - | Present | Cupredoxins | N/A | Non-toxic |
| **Aelin gp65** | - | - | - | Cytochrome C like | N/A | Non-toxic |
| **Aelin gp66** | - | - | - | TIM barrel | N/A | Non-toxic |
| **Aelin gp67** | Ribbon helix-helix DNA binding domain (1.7e-12) | - | - | DNA binding-3-helical bundle | Present | N/A |
| **Aelin gp68** | - | - | - | Belta grasp | N/A | Non-toxic |
| **Aelin gp70** | - | - | - | DNA binding-3-helical bundle | Present | Non-toxic |
| **Aelin gp71** | TPR repeat (5.7e-05) | - | - | Cytochrome C like | N/A | Non-toxic |
| **Aelin gp72** | CHAT domain (9.3e-06) | - | - | Belta grasp | N/A | Non-toxic |
| **Aelin gp73** | - | - | - | Cytochrome C like | N/A | Non-toxic |
| **Aelin gp75** | - | - | - | DNA binding-3-helical bundle | Present | Non-toxic |
| **Aelin gp76** | - | - | - | DNA binding-3-helical bundle | N/A | Non-toxic |
| **Aelin gp77** | - | - | - | Immunoglobulin-like | N/A | Non-toxic |
| **Aelin gp78** | - | - | - | Immunoglobulin-like | N/A | Non-toxic |

| | | | | | | |
|---|---|---|---|---|---|---|
| **Aelin gp79** | - | - | - | FAD/NAD binding motif | N/A | Non-toxic |
| **Aelin gp80** | - | - | - | DNA binding-3-helical bundle | Present | Non-toxic |
| **Aelin gp81** | - | - | - | OB-fold | N/A | Non-toxic |
| **Aelin gp82** | - | - | Present | Cytochrome C like | N/A | Non-toxic |
| **Aelin gp84** | - | - | - | DNA binding-3-helical bundle | Present | Non-toxic |
| **Aelin gp85** | - | - | - | Belta grasp | N/A | Non-toxic |
| **Aelin gp86** | - | - | | DNA binding-3-helical bundle | Present | Non-toxic |
| **Aelin gp87** | - | - | - | Immunoglobulin-like | N/A | Non-toxic |
| **Aelin gp88** | - | - | - | DNA binding-3-helical bundle | Present | Non-toxic |
| **Aelin gp89** | - | - | - | Immunoglobulin-like | N/A | Non-toxic |
| **Aelin gp90** | - | - | - | Cupredoxins | N/A | Non-toxic |
| **Aelin gp91** | - | - | - | DNA binding-3-helical bundle | Present | Non-toxic |
| **Aelin gp92** | - | - | - | Viral coat and capsid protein | N/A | Non-toxic |
| **Aelin gp93** | - | - | - | DNA binding-3-helical bundle | Present | Non-toxic |
| **Aelin gp94** | - | - | - | DNA binding-3-helical bundle | Present | Non-toxic |
| **Aelin gp95** | - | - | - | DNA binding-3-helical bundle | Present | Non-toxic |
| **Aelin gp96** | - | - | - | Immunoglobulin-like | N/A | Non-toxic |
| **Aelin gp97** | - | - | - | Viral coat and capsid protein | N/A | Non-toxic |
| **Aelin gp98** | - | - | Present | OB-fold | N/A | Non-toxic |

| | | | | | | |
|---|---|---|---|---|---|---|
| **Aelin gp99** | - | - | - | Flavodoxin-like | N/A | Non-toxic |
| **Aelin gp100** | - | - | - | Cytochrome C like | N/A | Non-toxic |
| **Aelin gp101** | - | - | - | Immunoglobulin-like | N/A | Non-toxic |
| **Childish gp52** | - | - | - | Tim barrel | N/A | Non-toxic |
| **Childish gp53** | - | - | - | Belta grasp | N/A | Non-toxic |
| **GeneCoco gp67** | - | - | - | Tim barrel | N/A | Non-toxic |
